# Partial ligand-receptor engagement yields functional bias at the human complement receptor, C5aR1

**DOI:** 10.1101/515700

**Authors:** Shubhi Pandey, Xaria X. Li, Ashish Srivastava, Mithu Baidya, Punita Kumari, Hemlata Dwivedi, Eshan Ghosh, Trent M. Woodruff, Arun K. Shukla

**Author notes:** equal contribution; SP, XL, AS. To whom correspondence should be addressed: Trent M. Woodruff or Arun K. Shukla.

## Abstract

The human complement component, C5a, binds two different seven transmembrane receptors termed as C5aR1 and C5aR2. C5aR1 is a prototypical G protein-coupled receptor that couples to Gαi sub-family of heterotrimeric G proteins and β-arrestins (βarr) following C5a stimulation. Peptide fragments derived from the carboxyl-terminus of C5a can still interact with the receptor, albeit with lower affinity, and can act as agonists or antagonists. However, whether such fragments might display ligand bias at C5aR1 remains unexplored. Here, we compare C5a and a modified C-terminal fragment of C5a, C5a^pep^, in terms of G protein coupling, βarr recruitment, endocytosis and ERK1/2 MAP kinase activation at the human C5aR1. We discover that C5a^pep^ acts as a full-agonist for G protein coupling, while only displaying partial agonism for βarr recruitment. We also observe that whilst C5a^pep^ is significantly less efficient in inducing C5aR1 endocytosis compared to C5a, it exhibits robust activation of ERK1/2 phosphorylation at levels similar to C5a. Interestingly, C5a^pep^ displays full-agonist efficacy with respect to inhibiting LPS induced IL-6 secretion in human macrophages, but its ability to induce human neutrophil migration is substantially lower compared to C5a. Taken together, our findings reveal ligand-bias at C5aR1, not only with respect to transducer-coupling and receptor trafficking but also in terms of cellular responses. Our findings therefore establish a framework to explore additional levels of biased signaling and biased ligands at C5aR1 with therapeutic potential. More generally, our findings may be extended to discover biased ligands for the broad sub-family of chemokine GPCRs which also interact with chemokine ligands through a biphasic mechanism.

## Introduction

The complement peptide C5a, a potent chemotactic agent and an anaphylatoxin, is one of the most critical components of the human complement system (1). C5a is a 74 amino acid long peptide that is generated upon the enzymatic cleavage of complement component C5 by C5-converatse. Abnormal levels of C5a and subsequent signaling triggered by it are crucial in a range of inflammatory disorders including sepsis, rheumatoid arthritis and psoriasis (1,2). C5a exerts its effects via two seven transmembrane receptors namely the C5aR1 and C5aR2 (also known as C5L2) (3). Of these, C5aR1 is a prototypical GPCR that is expressed in macrophages, neutrophils and endothelial cells (3). Upon binding of C5a, C5aR1 couples to Gαi sub-type of heterotrimeric G protein resulting in inhibition of cAMP levels and mobilization of intracellular Ca++ (3). Subsequently, C5a also triggers the phosphorylation of C5aR1 followed by recruitment of β-arrestins (βarrs) and receptor internalization (3).

Structurally, C5a harbors four different helices and connecting loops, and it is stabilized by the formation of three disulphide bonds (4). C5a interacts with C5aR1 through two distinct interfaces, one involving the core of C5a with the N-terminus of the receptor while the other involves the carboxyl-terminus of C5a with the extracellular side of the transmembrane helices of C5aR1 (Figure 1A) (5). It has been proposed that the structural determinants for high-affinity binding are provided by the first set of interaction while the second set of interaction is responsible for driving functional responses through the receptor (5). Peptides derived from the carboxyl-terminus of C5a can bind to C5aR1, albeit with much lower affinity compared to C5a, and they can also trigger functional responses (6,7). Whether such peptides may induce differential coupling of the two major signal transducers namely the G protein and βarrs, and thereby, may exhibit biased responses, remains completely unexplored.

**Figure 1.**
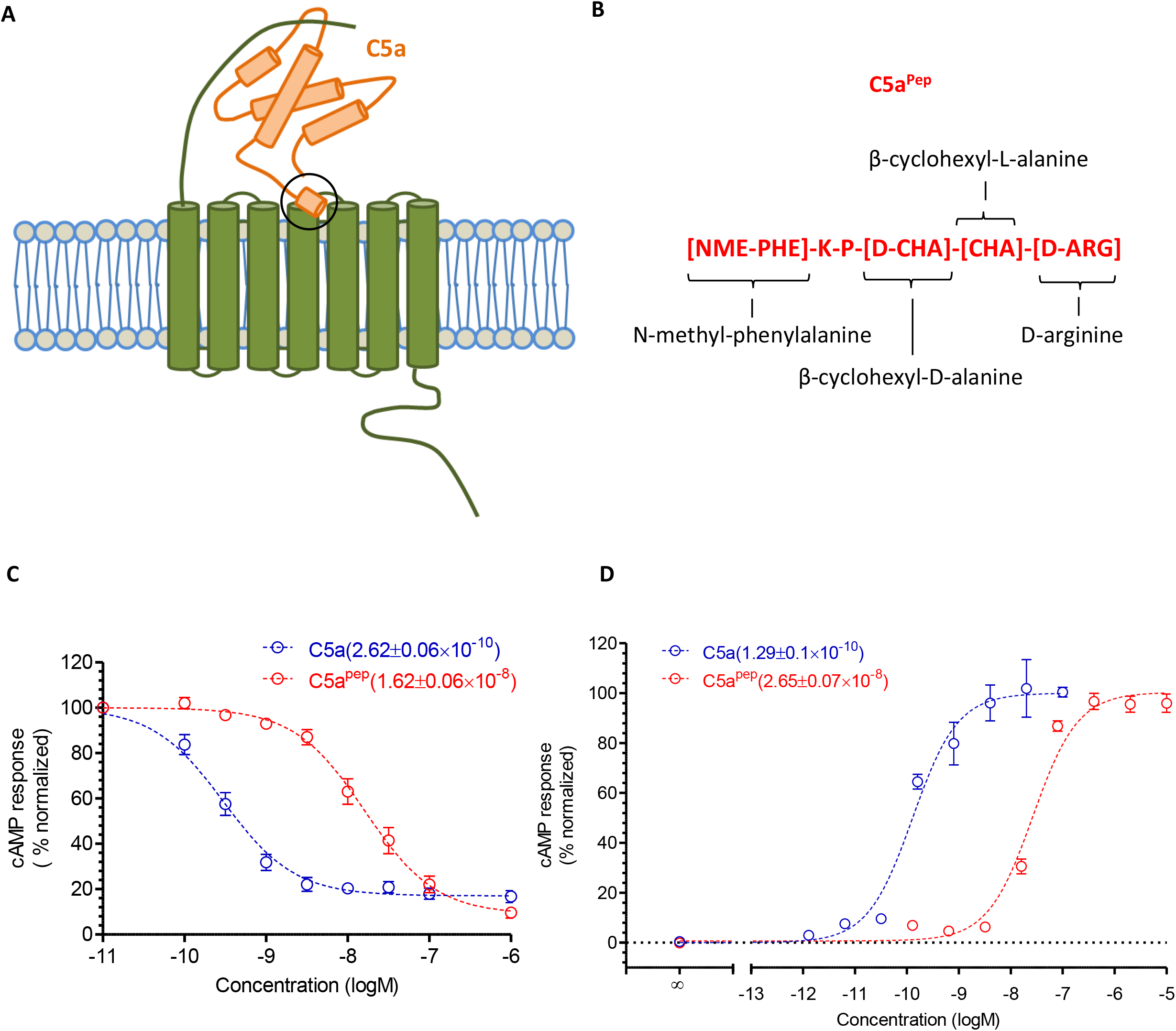
A modified carboxyl-terminus C5a peptide, C5a^pep^, is a full agonist for Gαi coupling. **A.** Schematic representation of C5a binding to C5aR1. There are two sites of interaction, one involving the N-terminus of C5aR1 and the other involving the extracellular loops. B. The primary sequence and modification of C5a^pep^, which is derived, based on the carboxyl-terminus of C5a. **C.** C5a^pep^ behaves as a full-agonist in GloSensor based cAMP assay. HEK-293 cells expressing C5aR1 were transfected with F22 plasmids. 24h post-transfection, the cells were stimulated with indicated concentrations of C5a and C5a^pep^ followed by recording of bioluminescence readout. **D.** Gαi-coupling induced by C5a and C5a^pep^ measured in CHO cells using LANCE cAMP assay. CHO cells stably expressing C5ar1 were stimulated with indicated concentrations of respective ligands. Data presented in panels C and D represent mean ± SEM of 3-5 independent experiments.

Here, we focus on a modified hexa-peptide, referred to as C5a^pep^ hereafter (Figure 1B), which displays highest binding affinity to C5aR1 amongst various C5a fragments (7), and characterize it vis-à-vis C5a with respect to transducer-coupling, functional outcomes and cellular responses. In particular, we measure the ability of C5a and C5a^pep^ to elicit Gαi-coupling, βarr recruitment and trafficking, receptor endocytosis, ERK1/2 MAP kinase activation, IL-6 release and neutrophil migration. We identify a significant bias in transducer-coupling, functional outcomes (i.e. endocytosis vs. ERK1/2 activation) and cellular responses (IL-6 release vs. neutrophil migration) between the two ligands which paves the way for subsequent characterization of physiological outcomes arising from biased signaling at C5aR1.

## Results and discussion

### C5a^pep^ is a full-agonist for Gαi-coupling

Although C5a^pep^ exhibits the highest binding affinity for C5aR1 amongst the peptides derived from, and modified based on, the carboxyl-terminus of C5a, its binding affinity for C5aR1 is still significantly lower than C5a (~70nM for C5a^pep^ and ~1nM for C5a) (7). We first measured the ability of C5a^pep^ to trigger Gαi-coupling to C5aR1 in HEK-293 cells using the GloSensor assay (8). Cells were stimulated with Forskolin to generate cAMP followed by incubation with various doses of C5a and C5a^pep^. We observed that both, C5a and C5a^pep^ inhibited cAMP level to a similar extent at saturating concentrations (Figure 1C) and with a similar time-kinetics (Figure S1). As expected, based on their binding affinities for the receptor, C5a^pep^ was approximately 100 fold less potent in cAMP inhibition compared to C5a (IC_50_ ~ 0.26 nM for C5a and IC_50_ ~ 16 nM for C5a^pep^). We also measured the efficacy of C5a^pep^ in C5aR1 expressing CHO cells using the LANCE cAMP assay (9) and observed a pattern of efficacy and potency very similar to that in HEK-293 cells (Figure 1D).

### C5a^pep^ is a partial-agonist for βarr recruitment

Upon C5a stimulation, C5aR1 undergoes phosphorylation and recruits βarrs which is important for receptor desensitization and internalization (10). Thus, we next measured the ability of C5a^pep^ to induce βarr coupling using a standard co-immunoprecipitation assay. There are two isoforms of βarrs known as βarr1 and 2 which exhibit a significant functional divergence despite a high-level of sequence and structural similarity (11). We expressed either βarr1 or βarr2 with FLAG tagged C5aR1 in HEK-293 cells and then measured their interaction upon ligand stimulation. As presented in Figure 2A, we observed a robust recruitment of both isoforms of βarrs upon stimulation of cells with C5a. Interestingly, the levels of βarr recruitment induced by C5a^pep^ were significantly lower compared to C5a, even at saturating ligand concentrations. As a control, we stimulated the cells with W54011, an antagonist of C5aR1 (12), and expectedly, it did not elicit any significant levels of βarr recruitment. In order to probe if there may be a temporal difference in C5aR1-βarr interaction pattern for C5a vs. C5a^pep^, we further carried out a time-course experiment for βarr recruitment. Still however, C5a^pep^-induced βarr recruitment was significantly lower than C5a (Figure 2B). Taken together with cAMP data presented above, these findings suggest that C5a^pep^ is biased towards Gαi-coupling over βarr recruitment as revealed by correlation plots using maximal responses in the two transducer-coupling assays (Figure 2C).

**Figure 2.**
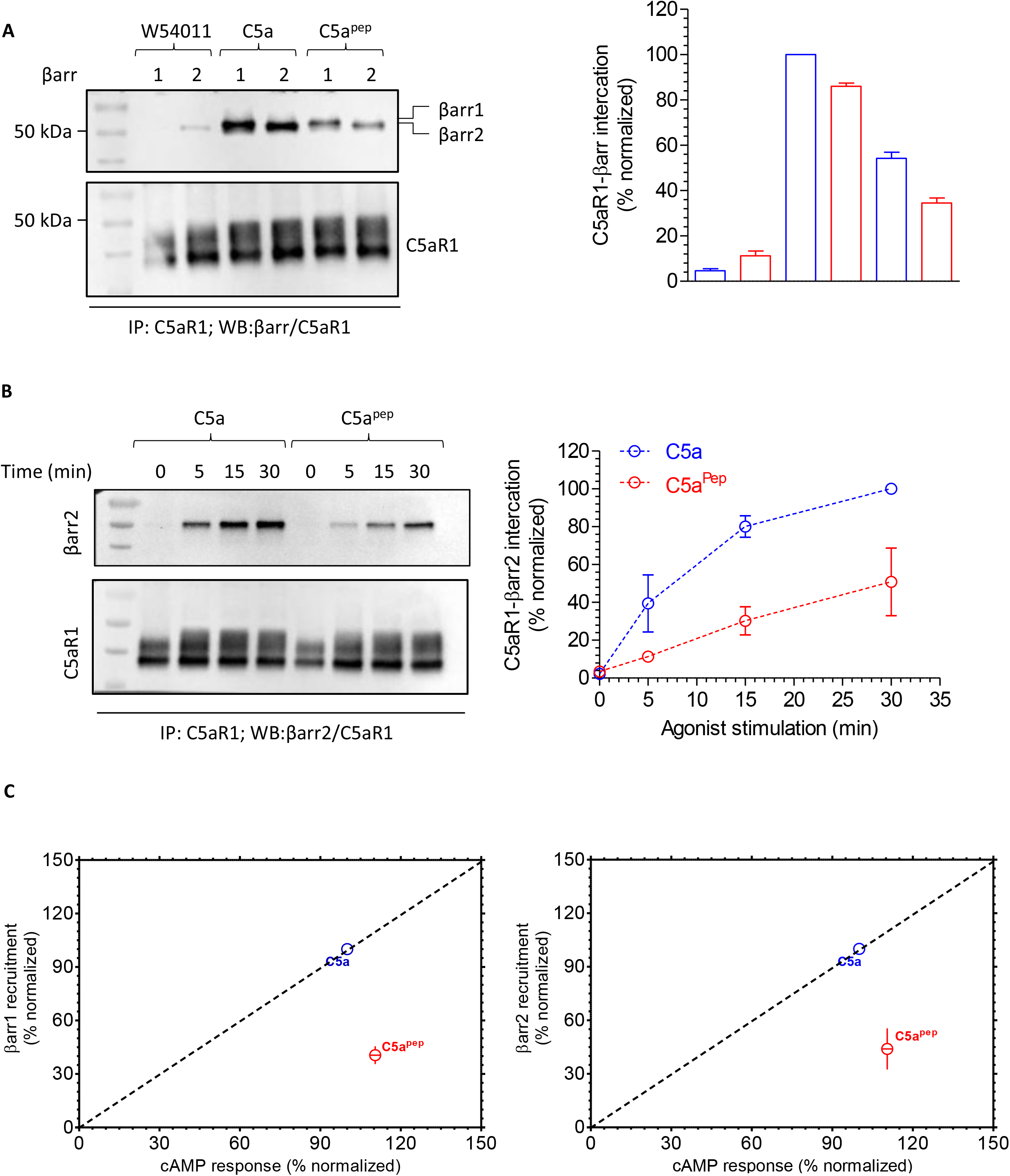
C5a^pep^ is a partial agonist for βarr recruitment. **A.** HEK-293 cells expressing Flag-C5aR1 and either βarr1 or 2 were stimulated with indicated concentrations of different ligands followed by crosslinking using DSP. Subsequently, Flag-C5aR1 was immunoprecipitated using anti-Flag antibody agarose and co-elution of βarrs were visualized using Western blotting. The right panel shows quantification of data. **B.** A time-course co-immunoprecipitation experiment to measure the interaction of C5aR1 with βarr2. The experiment was performed following the protocol as indicated in panel except that cells were stimulated for different time-points. The right panel shows densitometry-based quantification of the data (average ±SEM; n=3). **C.** Correlation plots derived using the maximal response in cAMP assay and βarr recruitment experiments presented above.

As mentioned earlier, C5a interacts with two 7TMRs, C5aR1 and C5aR2. Of these, C5aR2 (also known as C5L2) does not exhibit any detectable G-protein coupling although it robustly recruits βarrs upon agonist-stimulation (9,13,14). We observed that C5a^pep^ acts as a partial agonist for βarr2 recruitment, similar to C5aR1 (Figure S2). This finding suggests that the structural determinants of βarr recruitment in C5aR1 and C5aR2 are conserved while there must be a significant divergence at the level receptor conformation which dictates G-protein coupling.

### C5a^pep^ triggers slower endosomal trafficking of βarrs

βarrs are normally distributed in the cytoplasm and upon agonist-stimulation, they traffic to the membrane and interact with the receptors (15). Subsequently, upon prolonged agonist-stimulation, βarrs either dissociate from the receptor (class A pattern of βarr recruitment) or co-internalized with activated receptors in endosomal vesicles (class B pattern of βarr recruitment) (15). In order to probe whether C5a^pep^ might differ from C5a with respect to βarr trafficking patterns, we co-expressed C5aR1 with either βarr1-YFP or βarr2-YFP and visualized the trafficking of βarrs using confocal microscopy. We observed that C5a^pep^ was capable of promoting surface translocation of βarrs at similar levels as C5a during the early phase of agonist-stimulation (Figure 3, second panels). However, we observed that C5a^pep^ was significantly slower in promoting the appearance of βarrs in endosomal punctae and vesicles compared to C5a (Figure 3, third panels) although ultimately it did induce robust endosomal localization of βarrs (Figure 3, fourth panels).

**Figure 3.**
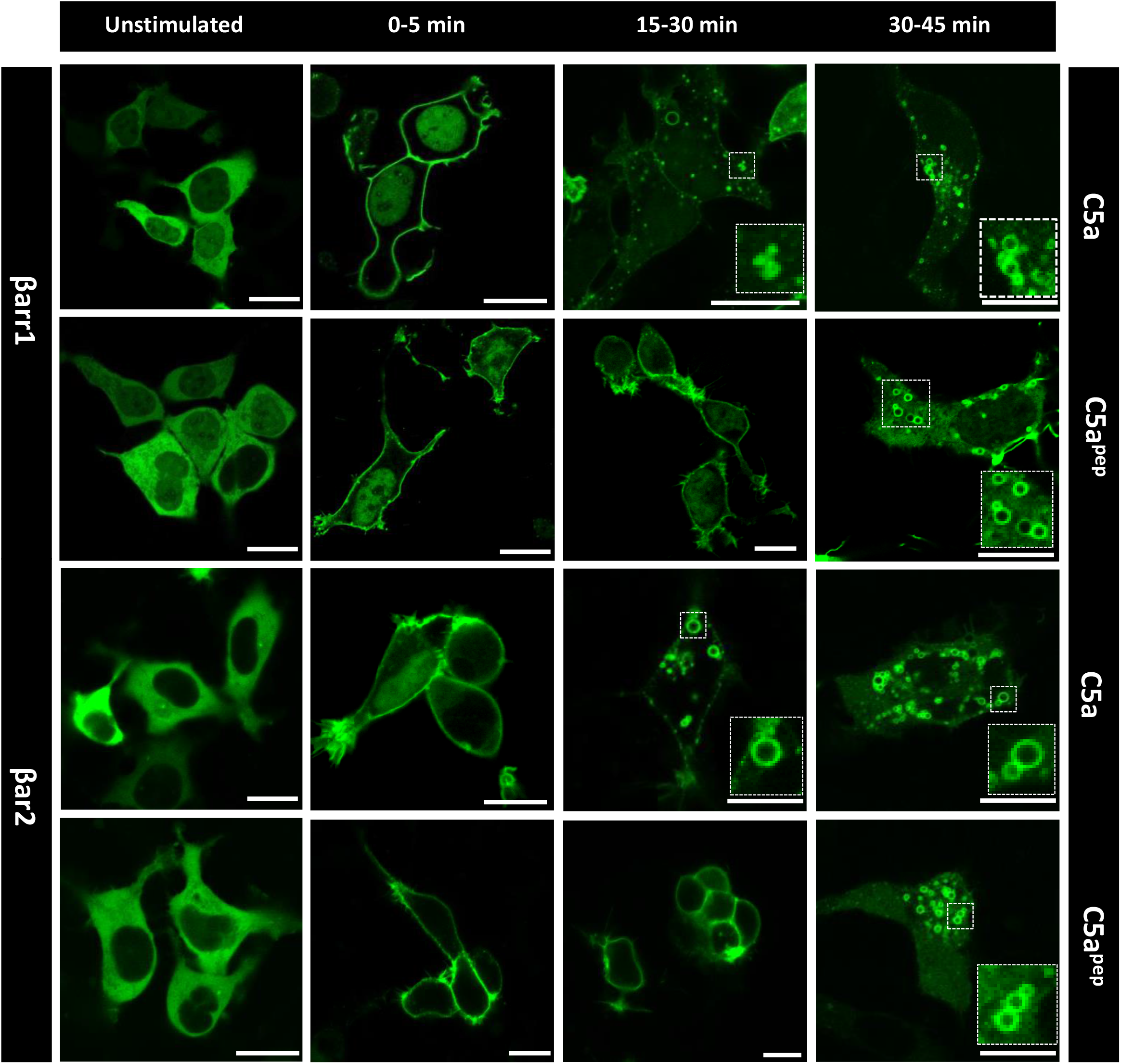
C5a^pep^ induces slower endosomal trafficking of βarrs. HEK-293 cells expressing C5aR1 and βarr1/2 were stimulated with C5a and C5a^pep^ and the trafficking patterns of βarrs were visualized using confocal microscopy for indicated time-points. Both, C5a and C5a^pep^ induced surface localization of βarrs at early time-points, however, somewhat slower endosomal trafficking of βarrs were observed for C5a^pep^ as assessed by localization of βarrs in endosomal vesicles. The figure shows representative images from three independent experiments.

### C5a^pep^ exhibits bias between ERK1/2 MAP kinase activation and receptor endocytosis

In order to probe whether bias in transducer coupling may translate into differential functional outcomes, we next measured the ability of C5a^pep^ to induce receptor endocytosis and ERK1/2 MAP kinase activation in HEK-293 cells. In agreement with our data on βarr recruitment and trafficking presented earlier in Figures 2 and 3, we observed a significantly lower levels of receptor endocytosis induced by C5a^pep^ in comparison to C5a (Figure 4A). This observation hints that βarrs may play a crucial role in endocytosis of C5aR1 and therefore, weaker βarr recruitment by C5a^pep^ translates into lower endocytosis. Interestingly however, C5a^pep^ was as efficacious as C5a in stimulating phosphorylation of ERK1/2 MAP kinase in HEK-293 cells, at least at the time-points that were tested in this experiment (Figure 4B-C). Thus, correlation of maximal levels of endocytosis triggered by C5a and C5a^pep^ with the ERK1/2 phosphorylation reveals a bias of C5a^pep^ in these two functional responses (Figure 4D).

**Figure 4.**
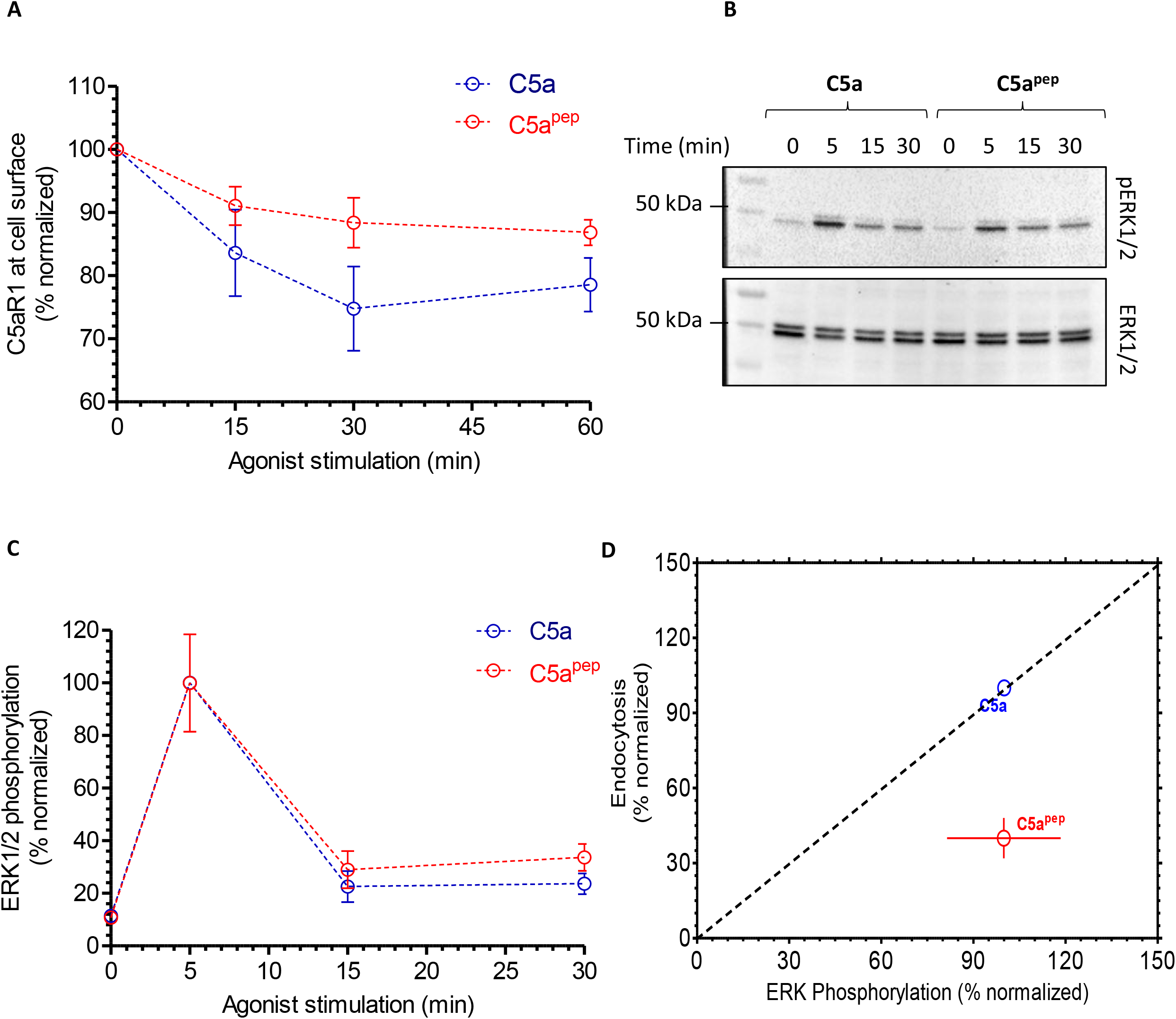
C5a^pep^ exhibits bias between receptor endocytosis ERK1/2 MAP kinase activation. **A.** HEK-293 cells expressing C5aR1 were stimulated with C5a and C5a^pep^ for indicated time-points followed by the assessment of surface receptor levels using a whole-cell ELISA assay. C5a^pep^ displays a weaker efficacy in promoting C5aR1 endocytosis compared to C5a. Data represent average±SEM from five independent experiments. **B.** C5a^pep^ induces robust activation of ERK1/2 MAP kinase at levels similar to C5a. HEK-293 cells expressing C5aR1 were stimulated with respective ligands for indicated time-points followed by measurement of ERK1/2 phosphorylation using Western blotting. C. Densitometry-based quantification of ERK1/2 phosphorylation data presented in panel B showing average ± mean of five independent experiments. **D.** Correlation plot derived using the responses of C5a and C5a^pep^ in endocytosis (at 30 min time-point) and ERK1/2 phosphorylation (at 5 min time-point).

### C5a^pep^ exhibits βarr isoform bias at a chimeric C5aR1

The interaction of βarrs with GPCRs is a biphasic process, which involves the receptor-tail (i.e. phosphorylated carboxyl-terminus) and the receptor-core (cytoplasmic surface of the transmembrane bundle) (16). In addition to the fully-engaged complexes involving both the tail and the core interaction, receptor-βarr complexes engaged only through receptor tail are functionally competent in terms of mediating receptor endocytosis and signaling (17–19). Based on the stability of their interaction with βarrs, GPCRs are categorized as class A and B which represent transient and stable interactions, respectively (15). Receptors having clusters of phosphorylatable residues in their carboxyl-terminus (such as the vasopressin receptor, V2R) typically interact stably with βarrs. As C5aR1 does not harbor such clusters, we generated a chimeric C5aR1 where we grafted the carboxyl-terminus of the V2R on C5aR1, and refer to this construct as C5a-V2R (Figure 5A). We first measured the cAMP response of C5aV2R upon stimulation with C5a and C5a^pep^, and observed a similar efficacy and potency profile as for the wild-type C5aR1 (Figure 5B). Interestingly however, we observed an equal recruitment of βarr1 by C5a^pep^ and C5a with C5a-V2R (Figure 5C). On the other hand, βarr2 recruitment induced by C5a^pep^ was still significantly weaker than C5a (Figure 5D). This resulted in a bias at the level of βarr isoform recruitment for the chimeric receptor as evident in the correlation plot (Figure 5E).

**Figure 5.**
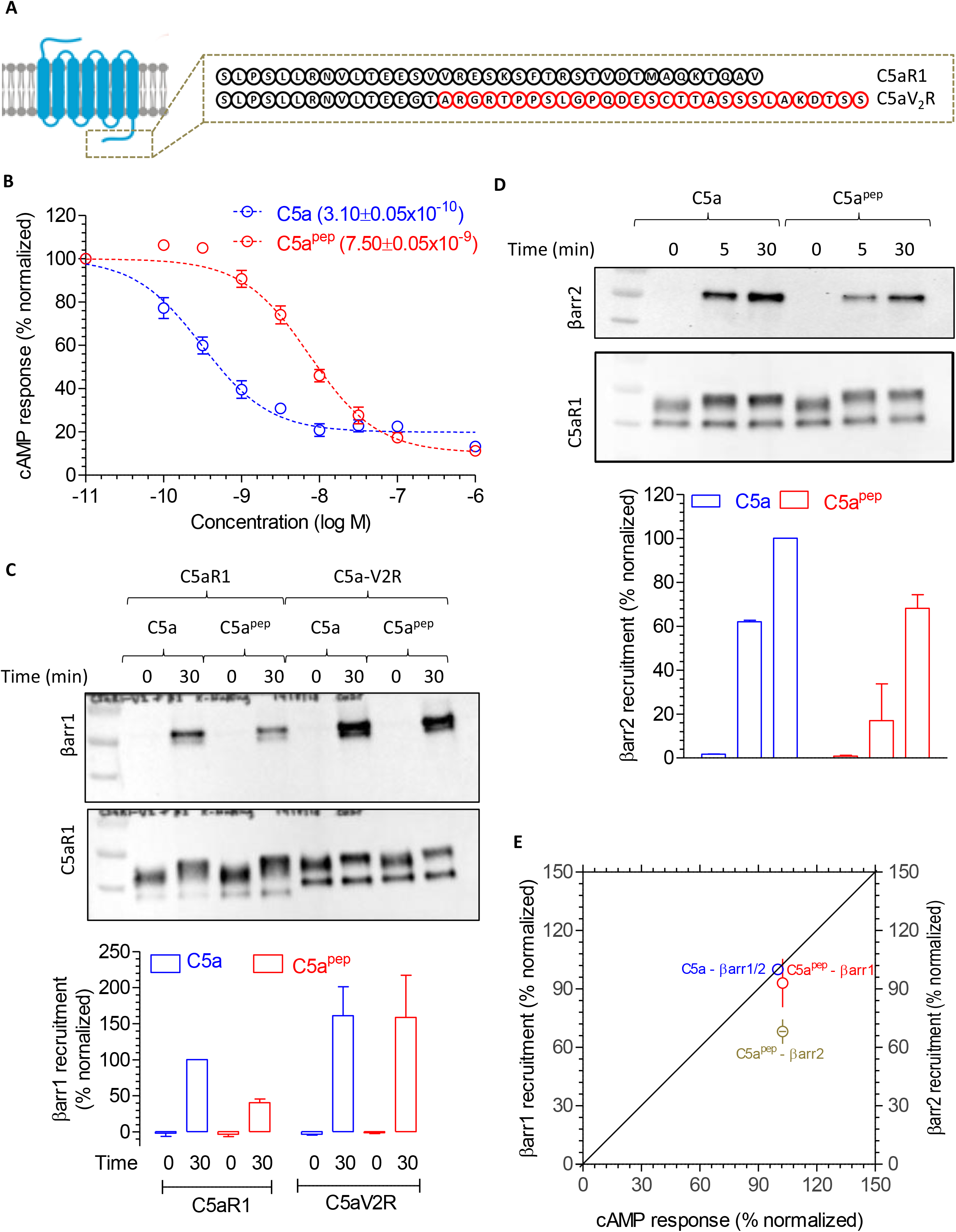
Transducer-coupling bias of C5a^pep^ is reversed at a chimeric C5aR1. **A.** Schematic representation of the carboxyl-terminus of C5aR1 and a chimeric construct harboring the carboxyl-terminus of AVPR2, referred to as C5aV2R. V2-tail in the chimeric construct is highlighted in red. **B.** C5a^pep^ behaves as a full-agonist in Glosensor based cAMP assay. HEK-293 cells expressing C5aV2R were transfected with F22 plasmids. 24h post-transfection, the cells were stimulated with indicated concentrations of C5a and C5a^pep^ followed by recording of bioluminescence readout. **C.** HEK-293 cells expressing Flag-C5aV2R and either βarr1 were stimulated with saturating concentrations of different ligands followed by cross-linking using DSP. Subsequently, Flag-C5aV2R was immunoprecipitated using anti-Flag antibody agarose and co-elution of βarr1 was visualized using Western blotting. The lower panel shows quantification of the data (average±SEM; n=3). **D.** Interaction of C5aV2R with βarr2 measured essentially in similar fashion as described in panel B (average ±SEM; n=3).E. Correlation plot of transducer-coupling derived using the maximal response in cAMP assay and βarr recruitment experiments presented above.

Our observation of βarr isoform preference for the chimeric receptor is important because several high-throughput screening assays, for example, Tango and enzyme-complementation, utilize chimeric GPCRs with V2R tail. This may result in miscalculation of ligand-bias and therefore, our data suggest that the patterns observed using chimeric receptors should be reanalyzed using wild-type receptors.

### C5a^pep^ elicits biased cellular responses

C5aR1 is endogenously expressed at high levels in macrophages and neutrophils, where it modulates multiple inflammatory responses (3). Stimulation of C5aR1 in human macrophages reduces LPS-induced release of IL-6 (2), whereas neutrophil C5aR1 activation induces rapid chemotaxis (3). To assess whether C5a^pep^ might exhibit a bias at the level of these cellular responses, we utilized HMDMs and PMNs to measure IL-6 release and migration, respectively. First, we measured Ca++ mobilization in HMDMs upon stimulation by C5a and C5a^pep^ and observed, once again, a full-agonist profile of C5a^pep^ (Figure 6A and S3). In addition, the inhibition of LPS-induced IL-6 release in HMDMs was also comparable for both C5a and C5a^pep^ (Figure 6B). Notably however, we observed that C5a^pep^ displayed a significantly blunted response in neutrophil migration compared to C5a even at saturating doses (Figure 6C). These observations therefore reveal a bias at the level of cellular responses exhibited by C5a^pep^ as obvious in the correlation plot (Figure 6D). These findings suggest that the bias displayed by C5a^pep^ at the level of transducer-coupling and functional responses measured in HEK-293 cells is preserved and translated into a cellular response bias in primary cells at the endogenous level of receptor.

**Figure 6.**
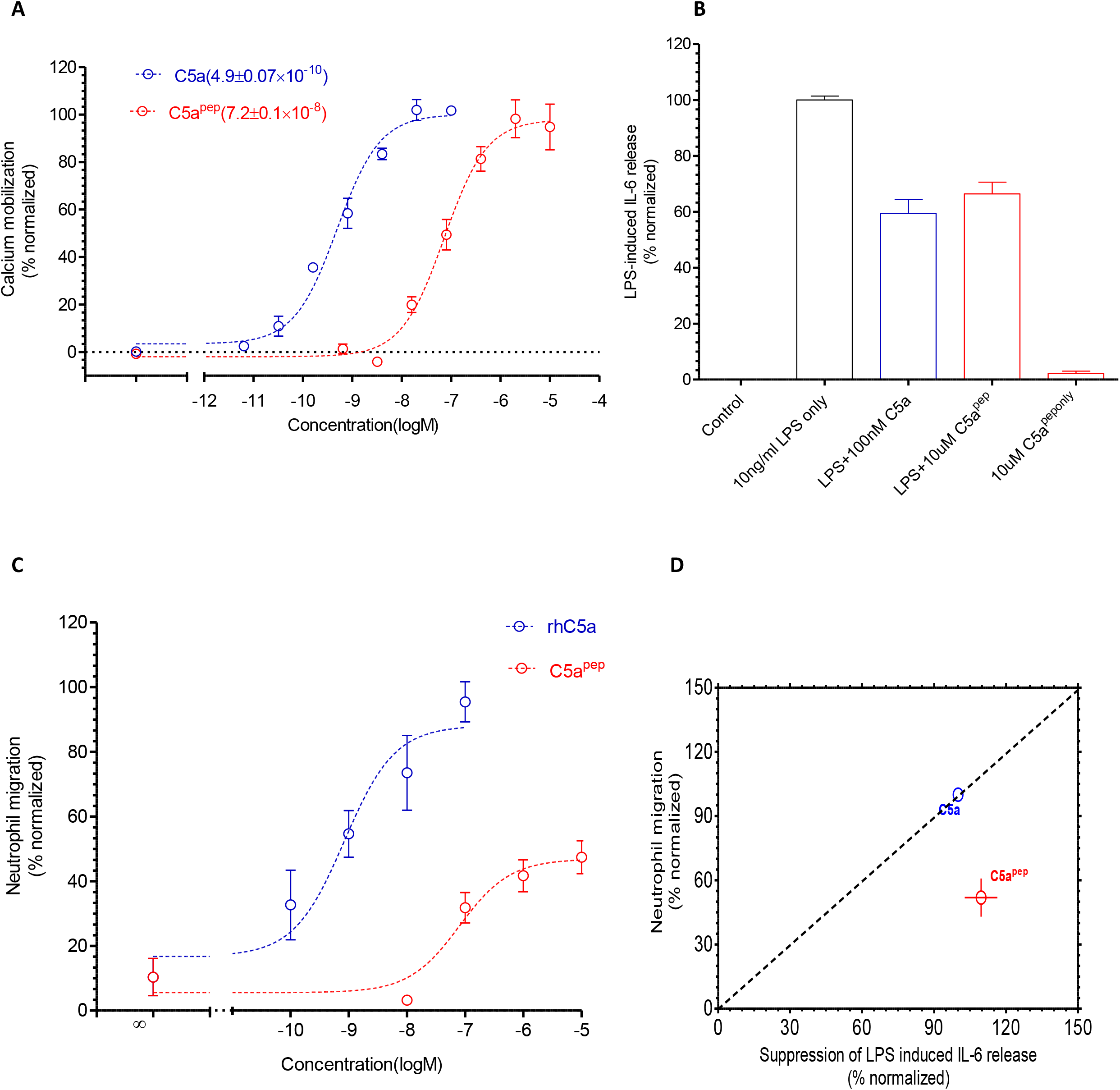
C5a^pep^ exhibits bias at the level of cellular responses. **A.** C5a^pep^ is a full-agonist in Ca++ mobilization assay in HMDM cells. HMDMs were first loaded with Fluo-4 calcium indicator followed by addition of respective ligands and subsequent measurement of fluorescence intensity. Data were normalized with maximal response obtained for C5a and represent mean ± SEM of triplicate experiments performed from three independent donors. **B.** C5a^pep^ behaves as a full-agonist for lowering of LPS-induced IL-6 release in HMDM cells. HMDMs were stimulated using 10 ng/ml LPS in the absence or presence of C5a or C5a^pep^. Subsequently, the levels of IL-6 present in the supernatant after 24h stimulation were quantified using ELISA, background-corrected with the values obtained for SFM/BSA and normalized with maximal response for LPS (i.e. treated as 100 %). Data represent mean±SEM of triplicate measurements conducted in cells from four independent donors. **C.** C5a^pep^ elicits only partial-agonist response in migration of human PMNs. Freshly isolated PMNs seeded into trans-well inserts were stimulated with respective ligands added to the receiver wells and then allowed to migrate for 1h. The number of migrated cells was recorded and normalized to the maximal C5a-induced migration. Data represents mean ± SEM of triplicate measurements conducted in cells from 3 independent donors. **D.** The correlation plot between the LPA-induced IL-6 release and PMN migration reveals bias induced by Ca5^pep^ at the level of cellular responses.

Interestingly, C5a^pep^ stimulation resulted in only a sub-maximal phosphorylation of ERK1/2 in HMDMs compared to C5a (Figure S4). This is in contrast with our observation in HEK-293 cells mentioned above where C5a^pep^ stimulates ERK1/2 phosphorylation at levels similar to C5a (Figure 4B-C). This is particularly interesting considering that C5a^pep^ behaves as full-agonist in Ca++ mobilization experiment in HMDMs. There is growing evidence for context-specific effector-coupling and functional responses downstream of several GPCRs (20,21). These emerging findings have refined our current understanding of biased signaling by providing substantial evidence of additional levels of complexities in GPCR signaling. Our data with C5a^pep^, especially in the context of ERK1/2 activation in HMDMs, further adds to this important paradigm of biased GPCR signaling.

It is important to mention here that chemokines, like complement C5a, also interact with their cognate GPCRs through a biphasic mechanism. Thus, it is tempting to speculate that fragments derived from the C-terminus of chemokines may also exhibit biased signaling through their receptors. Our findings also suggest that a more comprehensive analysis of C5a fragments might yield βarr-biased ligands at C5aR1. Although crystal structures of C5aR1 bound to small molecule antagonists have been determined recently (22,23), C5a-bound structure is still not available. High-resolution structure details of C5a-bound C5aR1 in future may provide structural insights into differential engagement of C5a^pep^ compared to C5a and how these differential interactions in the ligand-binding pocket yield transducer-coupling bias.

In summary, we discover C5a^pep^ as a biased C5aR1 agonist at the levels of transducer-coupling, functional outcomes and cellular responses. Going forward, an interesting avenue might be to evaluate the physiological responses elicited by C5a^pep^ *in-vivo.* Given that C5a attenuates LPS-mediated cytokine production from macrophages ((24) (Fig 6), a biased ligand such as C5a^pep^ which retains this beneficial activity, whilst diminishing the more pro-inflammatory activities of neutrophil migration, may be a novel therapeutic approach for inflammatory disorders. Furthermore, as we have demonstrated the recruitment of both isoforms of βarr to C5aR1, it would be interesting to evaluate the shared and distinct roles of these βarr in regulation and signaling of C5aR1. It is also notable that the second receptor activated by C5a (C5aR2), does not exhibit any detectable G protein coupling but displays robust βarr recruitment (13). Thus, C5aR2 may represent a natural example of a βarr-biased receptor making it an attractive target to further investigate specific role of βarrs in C5a mediate signaling.

## Materials and methods

### General reagents, constructs and cell culture

Most of the reagents were purchased from Sigma unless mentioned otherwise. Coding region of human C5aR1 was cloned in pcDNA3.1 vector with N-terminal signal sequence and FLAG-tag. For the construction of chimeric C5a-V2R construct, human C5aR1(1–326) was amplified and cloned in a modified pcDNA3.1 vector containing coding region of V2R C-terminus along with N-terminal signal sequence and FLAG tag. Coding regions of bovine βarr1 and βarr2 were cloned in pCMV6-AC-mYFP vector with stop codon being placed before mYFP. C5ap^pep^ was synthesized from Genscript. HEK-293 cells (ATCC) were maintained in DMEM at 5% CO2. Recombinant human C5a (C5a) was either purchased from Sino Biological or purified following a previously published protocol using a plasmid kindly provided by Prof. Gregers Rom Andersen (Aarhus University) (4). Ultrapure lipopolysaccharide from *Escherichia coli* K12 strain was purchased from Invivogen. Bovine serum albumen (BSA) was purchased from Sigma. For cell culture, trypsin-EDTA, Hank’s Balanced Salt Solution (HBSS), HEPES, Dulbecco’s Modified Eagle’s Medium (DMEM), phenol-red free DMEM, Ham’s F12, Iscove’s Modified Dulbecco’s Medium (IMDM) and Penicillin-Streptomycin were purchased from Themo Fisher Scientific. Dulbecco’s phosphate-buffered saline (DPBS) was purchased from Lonza.

The following cell lines were cultured as previously described (25). Chinese hamster ovary cells stably expressing the human C5aR1 (CHO-C5aR1) were maintained in Ham’s F12 medium containing 10% foetal bovine serum (FBS), 100 U/ml penicillin, 100 μg/ml streptomycin and 400 μg/ml G418 (Invivogen). Human embryonic kidney-293 (HEK293) cells were maintained in DMEM medium containing 10% FBS, 100 IU/ml penicillin and 100 μg/ml streptomycin. To ensure the consistency of cell function, cell morphology was continually monitored and neither cell line was used beyond passage twenty.

To generate human monocyte-derived macrophages (HMDM), human buffy coat blood from anonymous healthy donors was obtained through the Australian Red Cross Blood Service. Human CD14+ monocytes were isolated from blood using Lymphoprep density centrifugation (STEMCELL) followed by CD14+ MACS magnetic bead separation (Miltenyi Biotec). The isolated monocytes were differentiated for 7 days in IMDM supplemented with 10% FBS, 100 U/ml penicillin, 100 μg/ml streptomycin and 15 ng/ml recombinant human macrophage colony stimulating factor (hM-CSF) (Peprotech) on 10 mm square dishes (Sterilin). Non-adherent cells were removed by washing with DPBS and the adherent differentiated HMDMs were harvested by gentle scraping.

Human peripheral blood neutrophils (hPMN) were obtained from venous whole blood (20 ml) collected from healthy volunteers under informed consent. Samples were collected using venepuncture into BD K2EDTA Vacutainer^®^ blood collection tubes and processed within 5 hours. For neutrophil isolation, the anticoagulated blood was firstly layered over a Lymphoprep (STEMCELL) density gradient and centrifuged (800×g, 30 min, 22°C). The cell pellet in the densest layer of the gradient, containing a mixture of hPMN and erythrocytes was collected. The residual erythrocytes were removed using hypotonic lysis. Isolated neutrophils were counted and resuspended in a HBSS-based migration buffer (containing calcium and magnesium, supplemented with 20 mM HEPES and 0.5 % BSA) at a desired concentration. Cell viability was ≥ 92 % as assessed by Trypan blue exclusion on a Bio-Rad TC20™ automated cell counter.

### Preparation of C5aR1 expressing stable HEK-293 cell line

50-60% confluent HEK293 cells were transfected with 7μg FLAG-tagged C5aR1 DNA complexed with 21μg PEI (polyethylenimine). Next day, stable selection was started with optimal dose of G418 along with untransfected cells kept as negative control. After completion of stable selection, clonal population was prepared by limited dilution method. Highest expressing clones were propagated further and kept under G418 selection throughout the course of experiments. Surface exressionof C5aR1 was measured using a previously described whole cell surface ELISA protocol (26).

### ERK1/2 phosphorylation assay

Agonist-induced ERK1/2 phosphorylation was measured primarily using a previously described Western blotting based protocol (27). C5aR1 expressing stable cell lines were seeded into 6-well plate at a density of 1 million cells per well. Cells were serum starved for 12 hour followed by stimulation with 1μM of C5a and 10μM of C5a^pep^ respectively at selected time points. After the completion of time course, media was aspirated and cells were lysed in 100μl 2x SDS dye per well. Cells were heated at 95°C for 15 minute followed by centrifugation at 15000 rpm for 10 minute. 10uL of lysate was loaded per well and separated in SDS-PAGE followed by western blotting. Blots were blocked in 5% BSA (in TBST) for 1 hour and incubated overnight with rabbit phospho ERK (cat #9101/CST) primary antibody at 1:5000 dilution. Blots were washed thrice with TBST for 10 minute each and incubated with anti-rabbit HRP-coupled secondary antibody (1:10000, cat#A00098/Genscript) for 1 hour. Blots were washed again with TBST for three times and developed with promega ECL solution on chemidoc (BioRad). Blots were stripped with low pH stripping buffer and then re-probed for total ERK using rabbit total ERK (cat#9102/CST) primary antibody at 1:5000 dilution.

The ligand-induced phospho-ERK1/2 signaling was assessed using the AlphaLISA *Surefire Ultra* p-ERK1/2 (Thr202/Tyr204) kit (PerkinElmer) following the manufacturer’s protocol. Briefly, HMDMs were seeded (50,000/well) in tissue culture-treated 96-well plates (Corning) for 24 hours and serum-starved overnight. All ligand dilutions were prepared in serum-free medium (SFM) containing 0.1% BSA. For stimulation, cells were incubated with respective ligands for 10 min at room temperature and then immediately lysed using AlphaLISA lysis buffer on a microplate shaker (450 rpm, 10 min). For the detection of phospho-ERK1/2 content, cell lysate (5 μl/well) was transferred to a 384-well ProxiPlate (PerkinElmer) and added to the donor and acceptor reaction mix (2.5 μl/ well, respectively) with 2-hour incubation at room temperature in the dark. On a Tecan Spark 20M, following laser irradiation of donor beads at 680 nm, the chemiluminescence of acceptor beads at 615 nm was recorded.

### Receptor Internalization Assay

50-60% confluent *HEK293* cells were transfected with 3.5μg of FLAG-tagged C5aR1 DNA by polyethylenimine method of transfection at DNA:PEI ratio of 1:3. 24 hour post-transfection, 0.15million cells per well were seeded in 24-well plate (pre-coated with 0.01% Poly-D-Lysine). After 24 hour, cells were serum starved for 6hour followed by stimulation with 1μM of C5a and 10μM C5a^pep^ respectively for selected time points. After stimulation, cells were washed once with ice-cold 1XTBS. Cells were then fixed with 4% (w/v) PFA on ice for 20min and washed thrice with TBS to remove PFA. Blocking was done with 1%BSA prepared in 1XTBS for 1.5 hour. This was followed by incubation of cells with HRP-conjugated anti-FLAG M2 antibody (Sigma) at a dilution of 1:2000 prepared in 1% BSA+1XTBS for 1.5 hour. Afterwards, cells were washed thrice with 1% BSA+1XTBS. Surface expression was measured by incubating cells with 200uL TMB (Genscript) per well and reaction was stopped by transferring 100uL of developed colored solution to a 96-well plate already containing 100uL of 1M H_2_SO_4_. Absorbance was read at 450nm in a multi-plate reader (Victor X4). For normalization, cell density was measured using janus green. Briefly, TMB was removed and cells were washed twice with 1XTBS followed by incubation with 0.2% (w/v) janus green for 15 min. Destaining was done with three washes of 1ml distilled water. Stain was eluted by adding 800uL of 0.5N HCl per well. 200uL of this solution was transferred to a 96-well plate and absorbance was read at 595nm. Data normalization was done by dividing A450 by A595 values.

### GloSensor Assay for cAMP Measurement

50-60% confluent *HEK293* cells were co-transfected with 3.5μg each of C5aR1 and 22F (promega) plasmids. 24 hour post-transfection, cells were trypsinized and harvested by centrifugation at 1500rpm for 10 minute. Media was aspirated and cells were resuspended in luciferin sodium solution (0.5mg/ml) (Gold Biotech) prepared in 1X Hanks Balanced Salt Solution (HBSS/Gibco) and 20mM HEPES pH 7.4. Cells were then seeded in a 96-well plate at a density of 0.4 million cells per well and kept at 37°C for 1.5 hour in the CO_2_ incubator followed by incubation at room temperature for 30 minute. Basal reading was read on luminescence mode of multi-plate reader (Victor X4) and cycles were adjusted until basal values were stabilized. Cells were then incubated with 1uM forskolin and readings were recorded until maximum luminescence values were obtained. This was followed by stimulation of cells with specified concentrations of C5a and C5a^pep^ and values were recorded for 1 hour. Data was normalized with respect to minimal stimulation dose of ligand after basal correction.

### Cross-linking and Co-immunoprecipitation (CoIP)

50-60% confluent *HEK293* cells were co-transfected with C5aR1 and βarr1/ βarr2 plasmids by PEI (as mentioned earlier). 48 hour post- transfection, cells were serum starved for 6 hours and stimulated with respective doses of C5a/C5a^Pep^, harvested and proceeded for cross-linking experiment. Cells were lysed by dounce homogenization in 20mM HEPES pH7.4, 100mM NaCl, 1X phosphatase inhibitor cocktail (Roche), 2mM benzamidine hydrochloride and 1mM PMSF. This was followed by addition of 1mM dithiobis(succinimidyl-propionate) from a freshly prepared 100mM stock in DMSO. Lysate was tumbled at room temperature for 40 minute and reaction was quenched by adding 1M Tris pH 8.5. Lysates were solublized in 1%(v/v) MNG for 1 hour at room temperature followed by centrifugation at 15000 rpm for 15min. Cleared supernatant was transferred to a separate tube already containing pre-equilibrated M1-FLAG beads supplemented with 2mM CaCl_2_. Solution was tumbled for 2hour at 4°C and washed alternately with low salt buffer(20mM HEPES pH 7.4, 150mM NaCl, 0.01% MNG, 2mM CaCl_2_) and high salt buffer(20mM HEPES pH7.4, 350mM NaCl, 0.01% MNG, 2mM CaCl_2_) respectively. The bound proteins were eluted in FLAG-elution buffer containing 20mM HEPES pH 7.4, 150mM NaCl, 2mM EDTA, 0.01% MNG and 250ug/mL FLAG peptide. Co-immunoprecipitated βarr was detected by western blotting using rabbit anti-βarr mAb (1:5000, CST Cat# D24H9). Blots were stripped and reprobed for receptor with HRP-conjugated anti-FLAG M2 antibody (1:5000). Blots were developed on Chemidoc (BioRad) and quantified using ImageLab software (BioRad).

### BRET assay for measuring the interaction of βarr2 with C5aR2

C5a-mediated βarr2 recruitment to C5aR2 was measured using bioluminescent resonance energy transfer (BRET) as previously described (9). Briefly, HEK293 cells were transiently transfected with C5aR2-Venus and βarr2-Rluc8 constructs using XTG9 (Roche). 24 hours post transfection, cells were gently detached using 0.05% trypsin-EDTA and seeded (100,000/well) onto white 96-well TC plates (Corning) in phenol-red free DMEM containing 5% FBS. On the following day, cells were firstly incubated with the substrate EnduRen (30 μM, Promega) for 2 hours (37 °C, 5 % CO2). On a Tecan Spark 20M microplate reader (37°C), the BRET light emissions (460-485 and 520-545 nm) were continuously monitored for 25 reads with respective ligands added after the first 5 reads. The ligand-induced BRET ratio was calculated by subtracting the Venus (520-545 nm) over Rluc8 (460-485 nm) emission ratio of the vehicle-treated wells from that of the ligand-treated wells.

### Confocal Microscopy

For visualization of ligand-induced βarr recruitment, *HEK293* cells were co-transfected with C5aR1 and βarr1-YFP or βarr2-YFP plasmids in 1:1 ratio (total 7μg) by PEI. 24 hour post-transfection, 1million cells were seeded in 35mm glass bottom dish pre-coated with 0.01% Poly-D-Lysine. After 24 hour, cells were serum starved for 6 hour and stimulated with respective doses of C5a and C5a^pep^. For live-cell imaging, images were acquired using Carl Zeiss LSM780NLO confocal microscope for specified time intervals and image processing was done in ZEN lite (Zen-blue/ ZEN-black) software from Zeiss.

### Intracellular calcium mobilization assays

Ligand-induced intracellular calcium mobilization was assessed using Fluo-4 NW Calcium Assay kit (Thermo Fisher Scientific) following the manufacturer’s instructions. Briefly, HMDMs were seeded (50,000/well) in black clear-bottom 96-well TC plates (Corning) for 24 hours before the assay. Cells were firstly stained with the Fluo-4 dye in assay buffer (1X HBSS, 20 mM HEPES) for 45 min (37 °C, 5 % CO_2_). Respective ligands were prepared in assay buffer containing 0.5 % BSA. On a Flexstation 3 platform, the fluorescence (Ex/Em: 494/516 nm) was continually monitored for a total of 100 seconds with ligand addition performed at 16 seconds.

### Chemotaxis assays

Ligand-induced hPMN migration was assessed using 6.5 mm Transwell polycarbonate membrane inserts with 3.0 μm pore (Corning) to create a modified Boyden chamber (28). Freshly isolated hPMNs were seeded onto inserts (500,000/well) for 20 min (37 °C, 5 % CO_2_) in a HBSS-based migration buffer as described in previous section. To initiate cell migration, respective ligands prepared in migration buffer were added to the receiver wells in duplicates. After 60-min migration (37 °C, 5 % CO2), the inserts were gently washed once with DPBS, and the residual cells on the upper side of the membrane were removed using a cotton swab. Migrated cells were detached by adding 500 μl/well Accumax solution (Invitrogen) to the receiver wells (10 min, room temperature) and then counted using a Bio-Rad TC20™ automated cell counter.

### Measurement of cytokines release using ELISA

The immunomodulatory effect of respective C5aR ligands on LPS-induced cytokine release was assessed in primary human macrophages as previously described (29). HMDMs were seeded in 96-well TC plates (100,000 /well) for 24 hours before treatment. All ligands were prepared in serum-free IMDM containing 0.1% BSA. For stimulation, cells were co-treated with LPS and respective C5aR ligands for 24 hours (37 °C, 5 % C0_2_). The supernatant was collected and stored at −20 *°C* till use. IL-6 levels in the supernatant were quantified using respective human enzyme-linked immunosorbent assay (ELISA) kits (BD OptEIA) as *per* the manufacturer’s protocol.

### Data collection, processing and analysis

All experiments were conducted in triplicate and repeated on at least 3 separate days (for cell lines) or using cells from at least 3 donors (for HMDMs) unless otherwise specified. Data was analyzed using GraphPad software (Prism 7.0) and expressed as mean + standard error of the mean (SEM). Data from each repeat was normalised accordingly before being combined. For all dose-response assays, logarithmic concentration-response curves were plotted using combined data and analyzed to determine the respective potency values.

## Acknowledgment

We thank the members of our laboratories for critical reading of the manuscript. The research program was supported by the DBT Wellcome Trust India Alliance (Intermediate Fellowship to A.K.S.—IA/I/14/1/501285), Department of Biotechnology, Government of India (Innovative Young Biotechnologist Award to A.K.S.—BT/08/IYBA/2014-3), LADY TATA Memorial Trust Young Researcher Award to A.K.S., Science and Engineering Research Board (SERB) (SB/SO/BB-121/2013), Council of Scientific and Industrial Research (CSIR) (37[1637]14/EMR-II), and the National Health and Medical Research Council of Australia (NHMRC; project grant 1082271). A.K.S. is an EMBO Young Investigator, and T.M.W. is supported by a NHMRC Career Development Fellowship (1105420). We thank Mr. Ravi Ranjan for helping with C5aR1 and C5aR1-V2R cloning and optimization of C5a purification.

## Author’s contribution

SP and AS performed most of the experiments in HEK-293 cells except the confocal microscopy which was carried out by MB and PK; HD assisted in GloSensor assay and characterization of C5aR1-V2R constructs. XL performed Ca++ response assay in CHO-K1 cells, IL-6 release and PMN migration experiments under the supervision of TW. All authors contributed manuscript writing and approved the final draft. AKS supervised the overall project.

---

**Figure S1.**
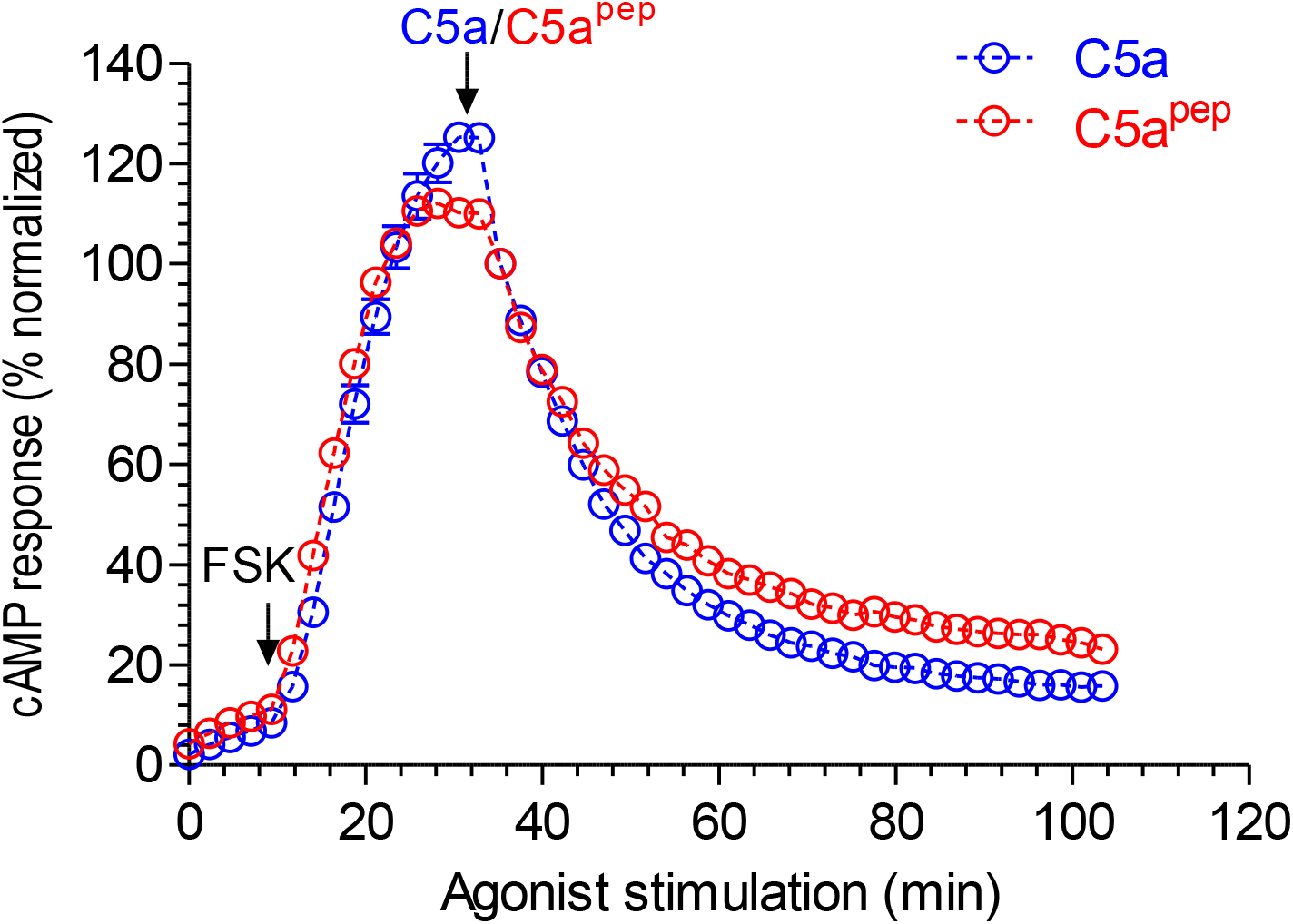
C5a^pep^ behaves as a full-agonist in GloSensor based cAMP time-kinetics assay. HEK-293 cells expressing C5aR1 were transfected with F22 plasmids. 24h post-transfection, the cells were stimulated with indicated concentrations of C5a and C5a^pep^ followed by recording of bioluminescence readout.

**Figure S2.**
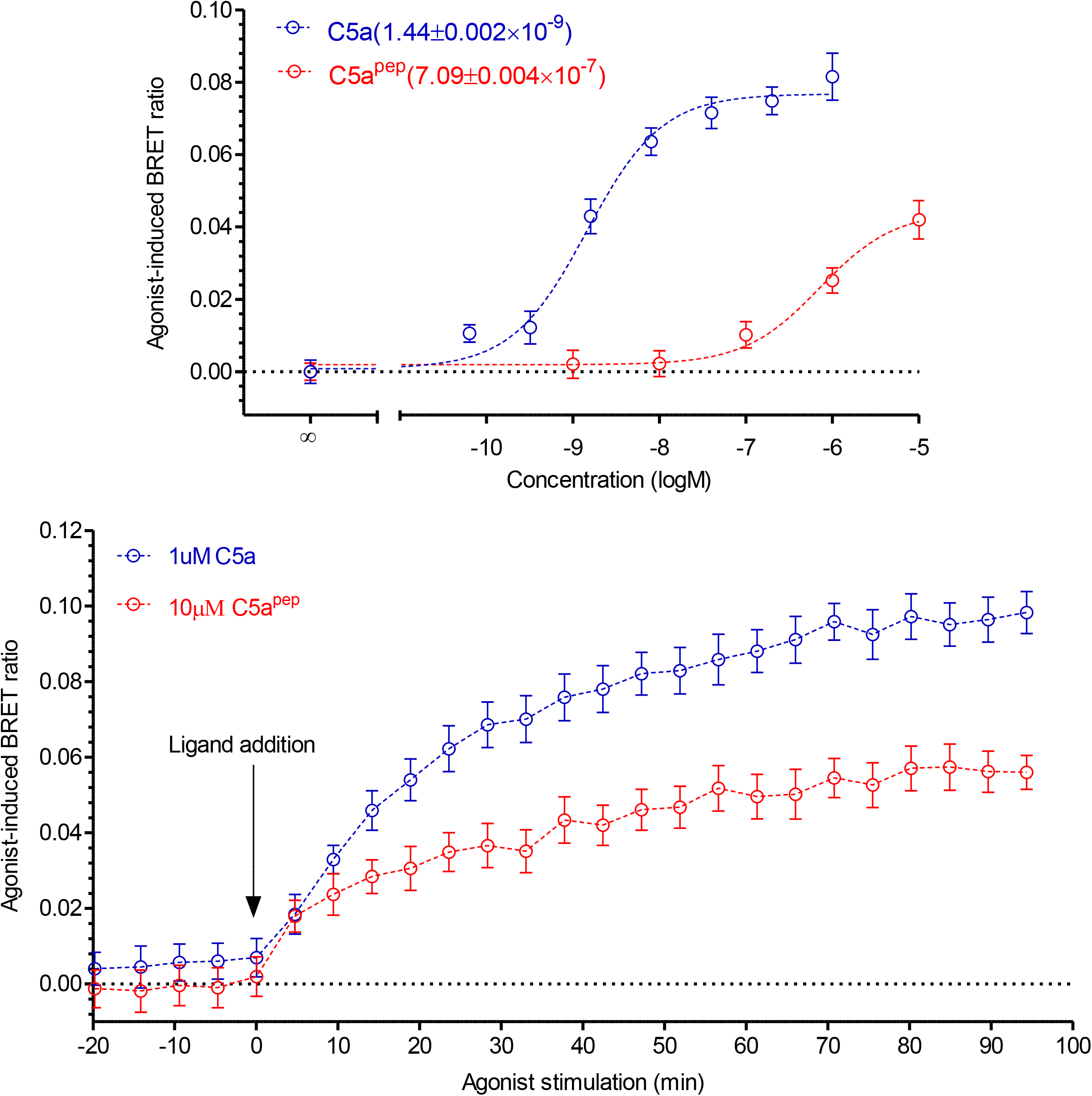
C5a^Pep^ promotes recruitment of βarr2 at C5aR2. HEK-293 cells expressing C5aR2-Venus and βarr2-Rluc8 constructs were firstly incubated with luciferase-substrate for 2h. Subsequently, cells were stimulated with respective ligands and BRET signals were monitored using either a dose response curve (upper panel) or time-kinetics (lower panel). Data represent average ± SEM of three independent experiments.

**Figure S3.**
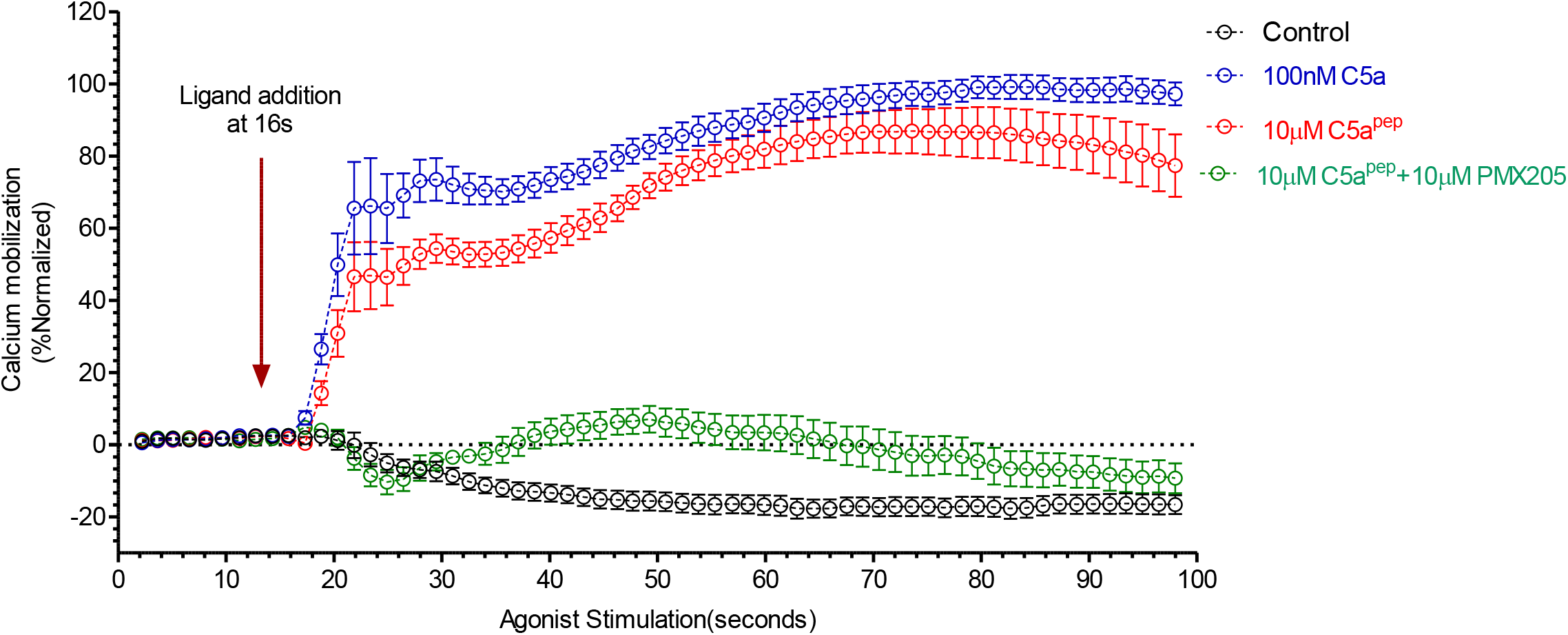
C5a^pep^ behaves as a full-agonist in Ca^++^ mobilization time-shift assay. HMDMs were stained with Fluo-4 dye in assay buffer followed by stimulation with respective ligands. Ligands were added at 16s and fluorescence was continually monitored for 100s.

**Figure S4.**
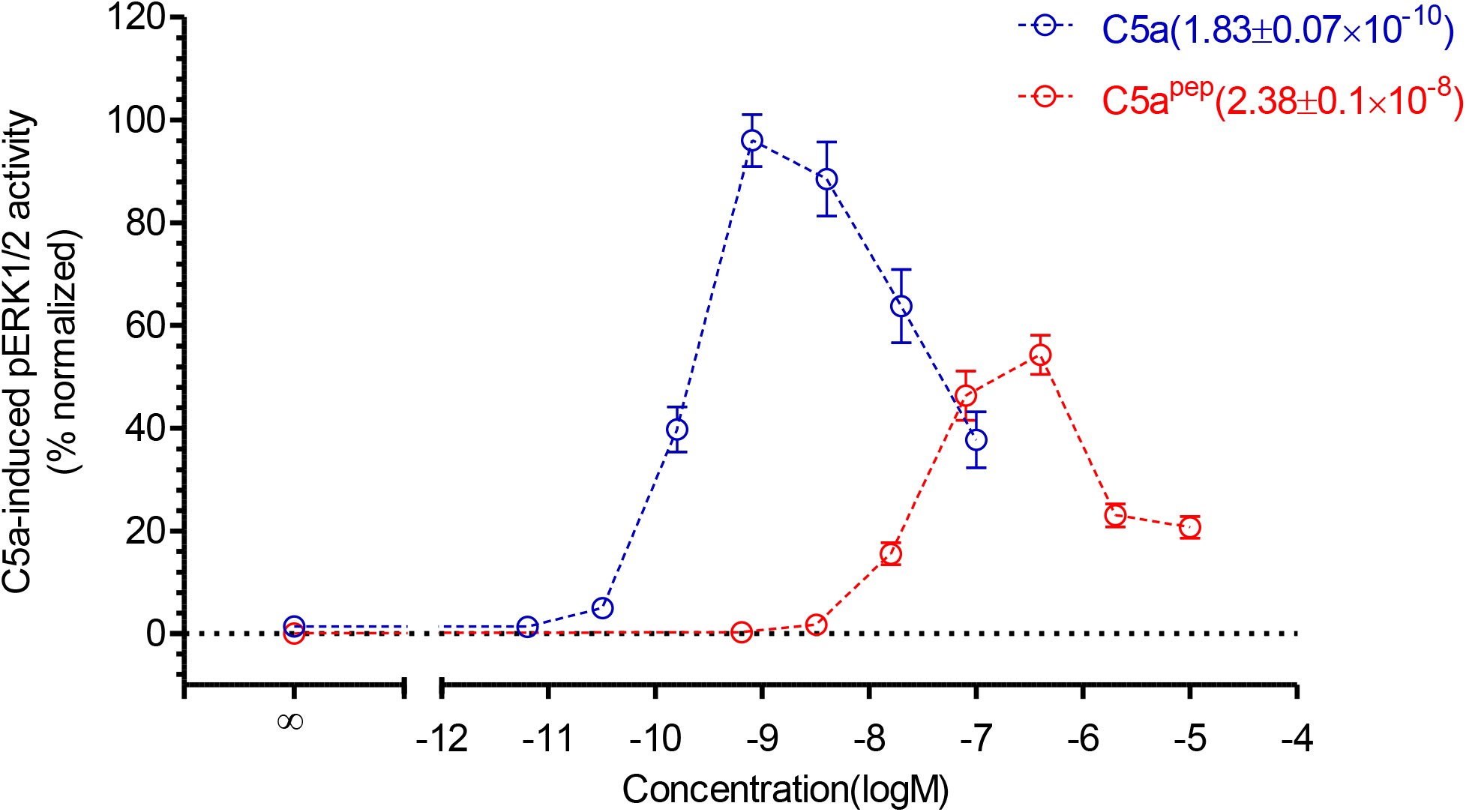
C5a^Pep^ induces sub-maximal ERK1/2 phosphorylation in HMDMs. HMDMs were serum starved overnight followed by incubation with respective ligands for 10min at room-temperature. Cells were lysed using AlphaLISA lysis buffer. For detection of phosphor-ERK1/2, cell lysate was added to donor and acceptor reaction mix and incubated at room temperature for 2hour. On a Tecan Spark 20M, following laser irradiation of donor beads at 680 nm, the chemiluminiscence of acceptor beads at 615 nm was recorded.

